# Identifying small proteins by ribosome profiling with stalled initiation complexes

**DOI:** 10.1101/511675

**Authors:** Jeremy Weaver, Fuad Mohammad, Allen R. Buskirk, Gisela Storz

**Affiliations:** Division of Molecular and Cellular Biology, *Eunice Kennedy Shriver* National Institute of Child Health and Human Development, Bethesda, MD 20892-5430, USA; Department of Molecular Biology and Genetics, Johns Hopkins School of Medicine, Baltimore, MD 21205, USA

**Keywords:** Small protein, alternate ORFs, leader peptide, antisense, genome annotation, Ribo-seq

## Abstract

Small proteins consisting of 50 or fewer amino acids have been identified as regulators of larger proteins in bacteria and eukaryotes. Despite the importance of these molecules, the true prevalence of small proteins remains unknown because conventional annotation pipelines usually exclude small open reading frames (smORFs). We previously identified several dozen small proteins in the model organism *Escherichia coli* using theoretical bioinformatic approaches based on sequence conservation and matches to canonical ribosome binding sites. Here, we present an empirical approach for discovering new proteins, taking advantage of recent advances in ribosome profiling in which antibiotics are used to trap newly-initiated 70S ribosomes at start codons. This approach led to the identification of many novel initiation sites in intergenic regions in *E. coli*. We tagged 41 smORFs on the chromosome and detected protein synthesis for all but three. The corresponding genes are not only intergenic, but are also found antisense to other genes, in operons, and overlapping other open reading frames (ORFs), some impacting the translation of larger downstream genes. These results demonstrate the utility of this method for identifying new genes, regardless of their genomic context.

**IMPORTANCE:** Proteins comprised of 50 or fewer amino acids have been shown to interact with and modulate the function of larger proteins in a range of organisms. Despite the possible importance of small proteins, the true prevalence and capabilities of these regulators remain unknown as the small size of the proteins places serious limitations on their identification, purification and characterization. Here, we present a ribosome profiling approach with stalled initiation complexes that led to the identification of 38 new small proteins.

## INTRODUCTION

Protein-protein interactions play an essential role in a variety of cellular processes, such as signal transduction and gene regulation. Small proteins, here considered to be 50 amino acids or fewer and encoded by small open reading frames (smORFs), have been shown to interact with and modulate the function of larger proteins (reviewed in (1-3)). These regulators have been identified in organisms spanning the phylogenetic tree of life, and important roles have been characterized for small proteins in bacteria and eukaryotes. In *Escherichia coli*, for example, the absence of the 49 amino acid protein AcrZ renders cells more susceptible to specific antibiotics (4), and cells lacking the 31 amino acid protein MgtS are sensitive to low magnesium concentrations (5, 6). In humans and other mammals, the small proteins myoregulin, sarcolipin, and phospholamban regulate muscle activity by affecting calcium transport (7, 8).

Despite the possible importance of small proteins, the true numbers of these regulators remain unknown as their small size limits their identification. ORF-finding algorithms traditionally employ a size limit for the scoring of genes (9) and apply a penalty for overlapping other ORFs (10). Their small size also often prevents these proteins from being accurately detected with protein gels, as they may run in the dye front and be poorly bound by SDS or protein dye (11). Traditional methods of purification are also biased against small proteins (12, 13), which have insufficient charge to bind ion-exchange columns and insufficient size to interact with non-reverse phase hydrophobic columns or be retained during dialysis. Additionally, small membrane proteins can bind nonspecifically to many column matrices due to their hydrophobicity. Finally, the few peptide fragments generated by proteolysis of small proteins limit detection by shot-gun proteomics (14). These challenges have stifled the detection of this class of proteins by standard methods.

As the importance of small proteins is being recognized, more focused searches for these proteins are being carried out (reviewed in (15)). Early genome-wide studies in *E. coli* utilized conservation of intergenic DNA sequences and the strength of ribosome-binding sites as a starting point for finding new small genes (16, 17). Similar approaches have been applied in eukaryotic organisms (18, 19), though the computational methods are more difficult, as both the increased size of the genome and the use of alternative splicing can mask small protein genes. In addition to the smORFs found in intergenic regions, there is growing recognition that transcripts can encode proteins in more than one ORF in the same region (reviewed in (20, 21)); these alternative ORFs (altORFs) generally code for smaller proteins than the originally annotated ORF, with some reported altORF-encoded proteins as small as 14 amino acids (22). Despite the success of computational methods in identifying new smORFs, it is likely that many small proteins have been missed (false negatives). Conversely it is critical that the synthesis of predicted small proteins be verified to avoid false positives.

Integration of data from large transcriptome analyses can improve the success of computational searches for smORFs. Ribosome profiling, a method that involves deep sequencing of ribosome-protected mRNA fragments, reveals the position of ribosomes throughout the transcriptome, clarifying which smORFs are translated under the conditions examined (reviewed in (23)). This approach has led to the identification of several small proteins (24-27), but again, there are limitations. Signals corresponding to altORFs encoded inside the confines of other genes can be swamped by the signal of the annotated gene. Another issue is that ribosome binding to an RNA does not prove that it leads to the production of a polypeptide (28). In eukaryotes, several signatures of profiling data that argue for translation are the presence of strong start and stop codon peaks as well as three nucleotide periodicity arising from the translocation of ribosomes one codon at a time. In bacteria, however, these signatures are weaker and more variable due to the lower resolution of the method, further complicating the discrimination of which transcripts are translated and which are merely ribosome-bound.

Although peaks in ribosome density at start and stop codons are the most useful in identifying new ORFs, the vast majority of ribosome-protected footprints in profiling data correspond to elongating ribosomes. In eukaryotes, the antibiotics harringtonine and lactimidomycin have been used to trap newly initiated 80S ribosomes at start codons and identify initiation sites (29, 30); elongating ribosomes are not inhibited by these antibiotics and continue elongation, terminating normally at stop codons. However, these compounds do not work in bacteria. Mori and co-workers found that treating *E. coli* cultures with tetracycline, an antibiotic that blocks tRNA binding in the ribosomal A site, leads to the accumulation of ribosome density at start codons. Using ribosome profiling of tetracycline-treated cells, they were able to re-annotate the N-termini of many known ORFs and discover candidate smORFs in intergenic regions (31). However, tetracycline traps ribosomes imperfectly at start codons. Only half the ribosomes on genes map to their start sites, blurring the signal. One promising alternative, Onc112, prevents initiation complexes from entering into the elongation phase (32, 33). Another promising substitute, retapamulin, a small molecule member of the pleuromutilin class, was previously shown to have a similar ability to specifically inhibit the first steps of elongation (34). The recent application of retapamulin in profiling experiments showed strong ribosome density at known start codons and little density attributable to elongating ribosomes; these data allowed the identification of start codons of altORFs within the coding sequences of other genes (35).

Here we present a strategy for identifying small protein genes in *E. coli* by combining traditional ribosome elongation data with information about initiation sites gleaned from profiling experiments conducted with retapamulin and Onc112. We sought to verify the synthesis of a subset of the predicted small proteins by assays to detect tagged derivatives and observed expression of 38 of the 41 genes tested. These results demonstrate that ribosome profiling with stalled initiation complexes provides a high confidence prediction of new small proteins in bacteria. Finally, the presence and location of these new smORFs reveals the density of information encoded by bacterial genomes.

## RESULTS

### Onc112 traps ribosomes at start codons but does not interfere with elongating ribosomes

The identification of initiation sites in eukaryotes has been aided by the use of antibiotics that enrich ribosome density at start codons in ribosome profiling experiments. Since such antibiotics have not been available for bacteria, we tested a promising candidate, Onc112, a proline-rich antimicrobial peptide (PrAMP) that binds in the exit tunnel and blocks aminoacyl-tRNA binding in the ribosomal A site (32, 36). Toeprinting analyses showed that Onc112 traps ribosomes at start codons, blocking elongation (33). We hypothesized that Onc112 should be selective for newly initiated 70S complexes because elongating ribosomes contain a nascent polypeptide that should prevent antibiotic binding. To test this possibility, we performed ribosome profiling on an untreated *E. coli* culture as well as one treated with 50 μM Onc112 for 10 min. As shown in Fig. 1a, ribosome density on the highly-expressed *lpp* gene is spread across the coding sequence in the untreated sample but is found almost exclusively at the start codon in the Onc112-treated sample. This effect holds genome-wide; in plots of ribosome density averaged over thousands of genes aligned at their start codons, a strong peak appears at the start codon while there is little or no density attributable to elongating ribosomes within coding sequences (Fig. 1b). These data show that like harringtonine and lactimidomycin in eukaryotes, Onc112 specifically traps ribosomes at start codons, while allowing elongating ribosomes to complete protein synthesis and terminate normally.

**FIG 1.**
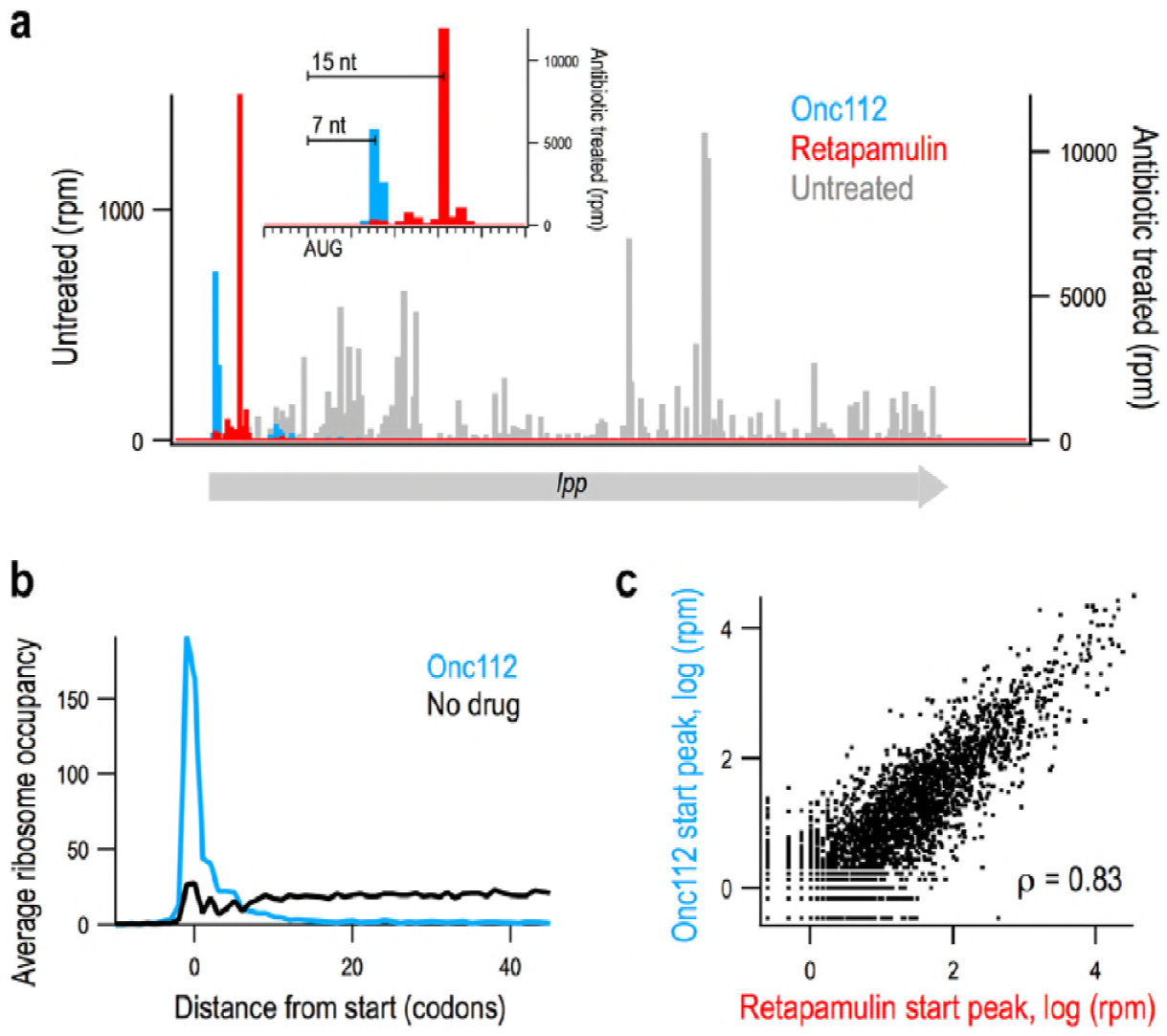
Onc112 and retapamulin similarly trap ribosomes at start codons. (a) Ribosome density on the *lpp* gene from Onc112-(blue) and retapamulin-treated (red) samples and an untreated control (grey). Inset: close-up view of the start site of *lpp*. (b) Average ribosome density at many genes aligned at their start sites in a sample treated with Onc112 and an untreated control. (c) Scatter plot of density at start sites in annotated genes in samples treated with Onc112 or retapamulin. The Spearman rank correlation is reported.

### Ribosome profiling signals for Onc112 and retapamulin are slightly different

A recent study used retapamulin and ribosome profiling to identify sites of non-canonical initiation within annotated ORFs (35). Like Onc112, retapamulin traps newly initiated 70S ribosomes at start codons while allowing elongating ribosomes to complete protein synthesis, such that ribosome density is strongly enriched at start codons (Fig. 1a). To compare the effects of Onc112 and retapamulin treatment, we calculated the intensity of start codon peaks on annotated ORFs, finding 3020 genes with any detectable ribosome density in the start codon region in both samples. There is a strong correlation between start peak intensity in these two antibiotic-treated samples (Spearman’s r = 0.83), arguing that both methods capture initiating ribosomes in a reproducible way (Fig. 1c).

Subtle differences may arise from variations in gene expression due to the different culture conditions; the retapamulin-treated sample was cultured in LB media whereas the Onc112-treated sample was obtained from a culture in complete synthetic MOPS media (Fig. S1). Another relevant difference in the sample preparation is that our protocol with Onc112 includes pelleting the sample over a sucrose cushion prior to nuclease treatment. This additional step depletes tRNAs from the ribosomal A site, allowing nucleases to cleave within the ribosome, thus shortening ribosome footprints. As a result, although the distance from the P-site codon to the 3’-boundary of the ribosome is reliably 15 nt in the retapamulin-treated library, it is more variable in our Onc112-treated library. Most often the peak is 6-10 nt downstream of the start codon; 7 nt in the *lpp* example (Fig. 1a). This difference is useful in annotating novel initiation sites: an AUG codon 6-10 nt upstream of density in the Onc112 data and 15 nt upstream of density in the retapamulin data has a high chance of being a bona fide start site and not a sequencing artifact.

### Onc112 and retapamulin can be used to identify putative translated smORFs

Given the ability of Onc112 and retapamulin to trap ribosomes at start codons at most annotated ORFs, we combined the information from these datasets to create a high-fidelity screening method for identifying new smORFs likely to be translated (Fig. 2a). We first generated a list of 160,995 smORFs of eight codons or longer whose start codons (AUG, GUG, or UUG) are 18 nt or more away from annotated coding regions (either protein coding sequences or functional RNA genes). We computed the ribosome density associated with each start site, including ribosome footprints 0-18 nt downstream of the first nt in the start codon. A CDF plot of these data are shown in Fig. 2b (left axis); the y-value reflects the percentage of predicted smORFs that have a start peak less than or equal to the x-value. This plot shows that ∼ 96% of the putative smORFs have no associated density at their start sites (x = 0). This means that ∼ 4% have start peaks greater than zero. Only 0.25% of the predicted smORFs had more than 5 reads per million mapped reads (rpm), as delineated by the dotted line in Fig. 2b. Thus, the vast majority of the putative smORFs likely are not translated.

**FIG 2.**
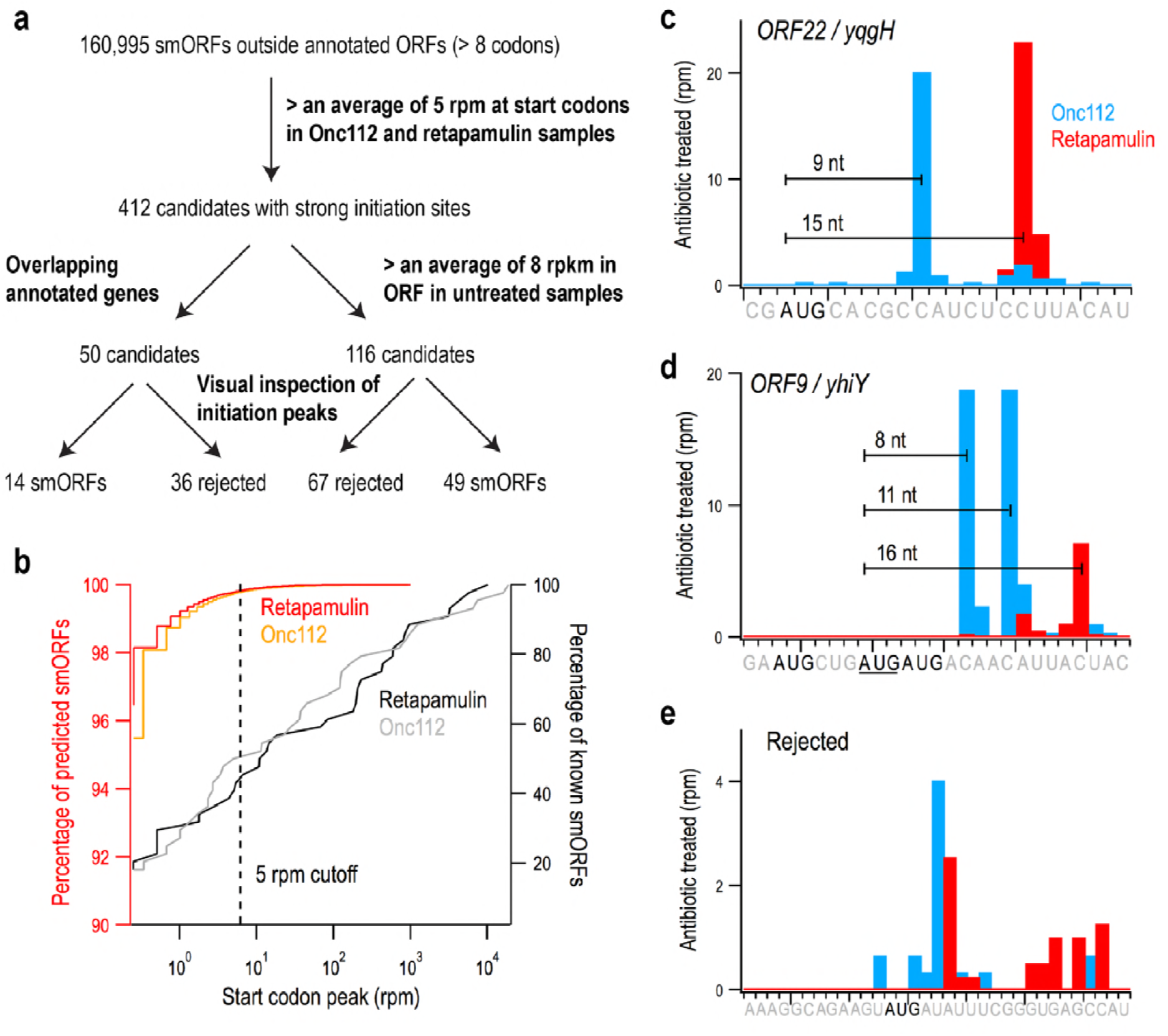
Using ribosome profiling data to discover new small open reading frames (smORFs). (a) Flow chart showing the criteria used to identify smORFs in intergenic regions. (b) CDF plot showing the percentage of known, annotated smORFs (n = 44, right axis, black and grey) on the y-axis less than or equal to the ribosome density near the start site (x-axis) as compared with candidate smORFs (n = 160,995, left axis, red and orange). Candidates with an average of > 5 rpm were selected for further screening (dotted line). (c) The proper spacing of ribosome density at start codons in treated samples helps to identify bona fide small protein coding genes such as ORF22 / *yqgH*. (d) In cases where several start codons could explain the ribosome density, spacing helps determine the correct site; ORF9 / *yhiY* likely initiates with the second AUG codon of the three shown. (e) Many candidates were rejected because the start site does not align properly with the density observed.

To calibrate our method for identifying new candidate smORFs, we examined the ribosome density on the start codons of smORFs previously shown to encode proteins. The test set included two different groups. The first group was comprised of 44 small proteins annotated initially together with small proteins identified by sequence conservation and strong matches to ribosome binding site models (16). The ribosome density after retapamulin or Onc112 treatment varied by four orders of magnitude (Table S1): ∼80% of this group had detectable signal at start sites and ∼60% of known smORFs had start peaks above 5 rpm (Fig. 2b, right axis). The second group of proteins had less conservation and weaker matches to ribosome binding site models but were shown to be synthesized as tagged derivatives in a recent study (17). Of the 36 proteins in the second set, ∼70% showed signal but only 20% had Onc112 or retapamulin reads above 5 rpm at the start site (Table S1), possibly due to the lower level of expression of these smORFs. Given that the majority of the small proteins in the first set of annotated smORFs have start peaks 5 rpm or higher (Fig 2b), we used this threshold to eliminate false positives in our list of putative smORFs; 412 novel smORFs above this threshold were selected for further consideration.

An important caveat in treating cells with Onc112 and retapamulin is that these antibiotics could enhance ribosome density on some initiation sites that are not normally used. The antibiotics dramatically increase the concentration of free 30S and 50S subunits given that they allow elongating ribosomes to complete protein synthesis and be recycled but block entry into the elongation cycle. The recycled subunits are free to initiate at less optimal start codons, where they will be trapped by the antibiotics. To remove these false positives, we used traditional ribosome profiling data (from untreated cells) to capture elongating ribosomes along the entire ORF. Of the 412 smORFs with strong start codon peaks, 116 had traditional ribosome profiling density above 8 reads per kilobase per million mapped reads (rpkm).

We next examined the 116 most promising candidates on a genome browser. In our screen for ribosome density at initiation sites (Fig. 2a), we summed the reads from 0-18 nt downstream of the first nucleotide in the start codon, an intentionally broad range. In our visual inspection, we searched for retapamulin peaks ∼15 nt and Onc112 peaks 6-10 nt downstream of the first nt in the start codon as seen for *lpp* (Fig. 1a). The same spacing is observed for the most promising candidates (e.g. *ORF22/yqgH* in Fig. 2c). In some cases of multiple possible start codons, we were able to readily predict the correct start based on distance (e.g. *ORF9/yhiY*, Fig. 2d). For most of the candidates that were rejected, the predicted start site did not align with the Onc112 or retapamulin ribosome density (Fig. 2e). Another source of false positives were smORFs close to highly-translated genes, such as ribosomal proteins, where the noise is high enough to pass the cutoff for start peaks and normal profiling density (data not shown). Based on these criteria, 67 candidates were rejected, leaving 49 candidates. Visual inspection proved helpful in refining the data, but in the future, our algorithms can be further developed to incorporate additional criteria for large-scale screens for candidate smORFs.

We also inspected 50 additional smORFs with strong start peaks (>5 rpm) for which we were unable to calculate rpkm values for elongating ribosomes because the smORFs overlap an annotated gene and the ribosome density cannot be assigned to one gene or the other. Upon inspection, 36 of these were rejected due to incorrect start site selection or high levels of noise due to adjacent highly-translated genes, leaving 14 of interest. In addition to these 14 candidates and the 49 discussed above, another three were discovered as the correct start sites for candidates that were rejected, and two more were discovered in a preliminary screen using similar cutoffs but a different collection of traditional ribosome profiling data.

Together, this workflow yielded 68 candidate smORFs with high start codon peaks and some level of traditional ribosome profiling data, including both independent genes and altORFs (Table S3). Initially, the smORFs were assigned numbers, but were renamed if we obtained evidence of small protein synthesis (see below). As expected, the majority of these candidates start with AUG codons (50), although GUG (9) and UUG (9) codons were also observed. A histogram of the predicted protein lengths is shown in Fig. S2: the majority of the predicted small proteins are 40 amino acids or shorter, although the analysis also identified seven candidates that were longer than 50 residues. A few of the candidates are overlapping in that they correspond to different possible start codons in the same frame.

### The majority of predicted small proteins are synthesized

To validate that the corresponding small proteins are synthesized, an SPA tag was integrated into the chromosome upstream of the stop codon of the 38 putative smORF genes with the highest ribosome density in the presence of the inhibitors and deemed the strongest candidates by the visual inspection. The tag allowed immunoblot analysis on the basis of the 3X FLAG epitope (Fig. 3). While the exposure needed to detect the small proteins varied significantly (as reflected in different levels of the background bands), 36 of the 38 tagged small proteins were detected in cells grown to exponential or stationary phase in LB at 37°C, conditions comparable to those used in the ribosome profiling experiments. The inability to detect the remaining two chromosomally-tagged smORFs (*ORF24* and *ORF56*) could stem from these smORFs being false positives in the screen or from the degradation of the tagged derivatives. We have nonetheless observed the expression the majority of the predicted genes, validating the predictive capability of utilizing multiple ribosome profiling datasets.

**FIG 3.**
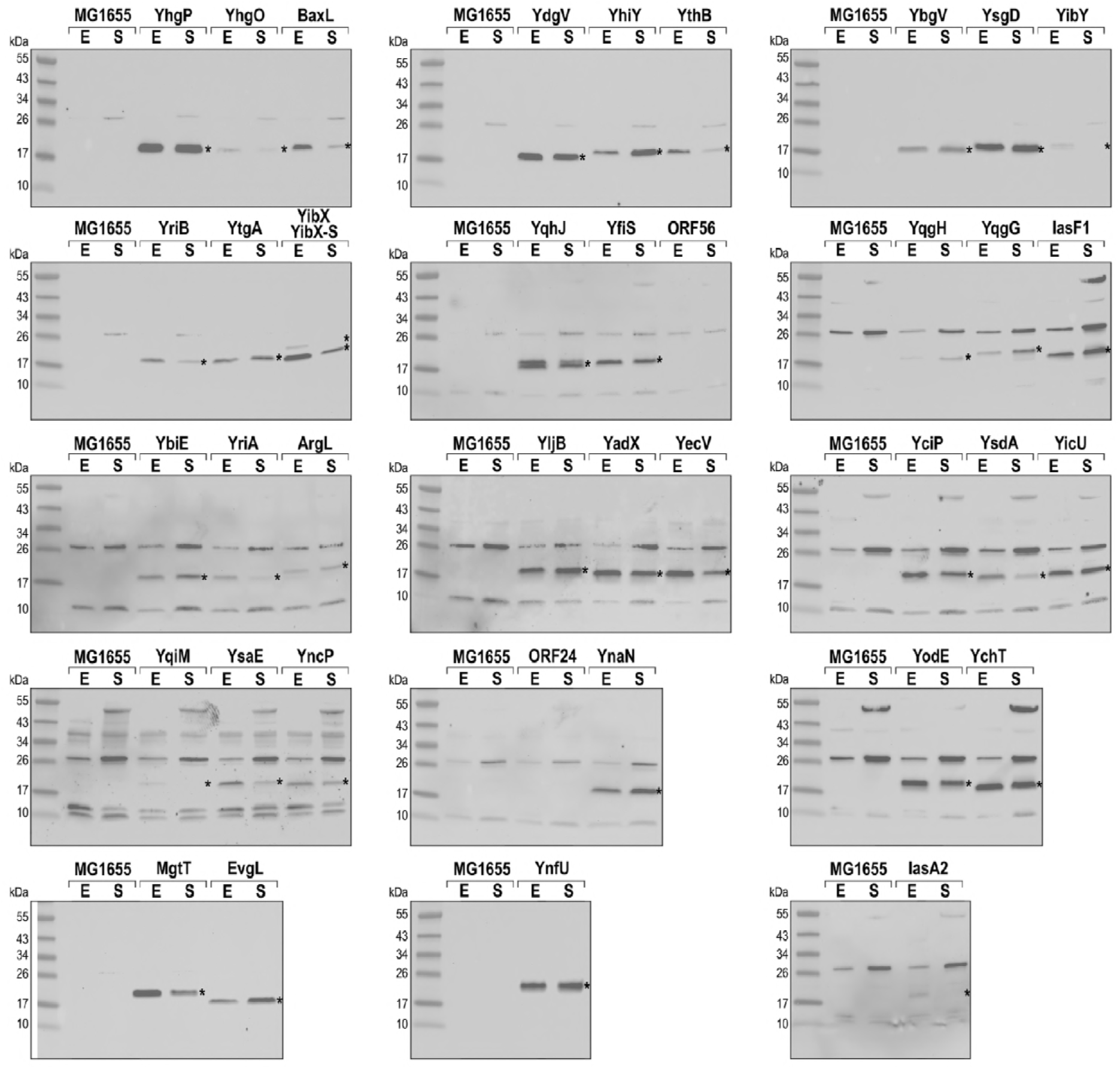
Western analysis confirms expression of 95% of tested predicted smORFs. MG1655 strains with chromosomally-tagged, putative smORFs were grown to exponential (E) and stationary (S) phase in rich media (LB). Gel samples were prepared to load equivalent numbers of cells based on OD_600_. Immunoblot analysis was conducted against the 3X FLAG motif included in the tag using HRP-conjugated, anti-FLAG antibodies. Wild-type MG1655 was included as a negative control. Blots requiring a longer exposure to show tagged proteins revealed more background bands. Bands corresponding to smORFs are marked with *.

Several previously detected small proteins are only expressed under very specific growth conditions (17, 37). As shown in Fig. 3, we observe that 10 newly-detected small proteins are present at >2-fold higher levels in exponential phase and four are present at >2-fold higher levels in stationary phase. The majority of the small proteins appear at roughly-equal levels during both of these growth phases but may be induced under other conditions.

### The levels of tagged small proteins span a wide range

As indicated above, the ability to detect the small proteins varied. To directly compare the overall levels of the proteins, both among themselves and with previously identified small proteins, we analyzed stationary phase samples of several examples of each group of proteins (Fig. 4). Among the newly-identified proteins, the levels of YnfU are highest, but these levels fall between the characterized multi-drug efflux pump regulator AcrZ (diluted 5-fold in Fig. 4) and uncharacterized protein YoaK, which, respectively, are among the better- and worse-expressed small proteins identified initially (16). The levels of the remaining small proteins cover a wide range, as is seen when comparing the samples (YsgD and YthB) loaded on two different gels as a reference. These blots also show that most of the other newly-identified small proteins are expressed at levels below YoaK under the conditions tested.

**FIG 4.**
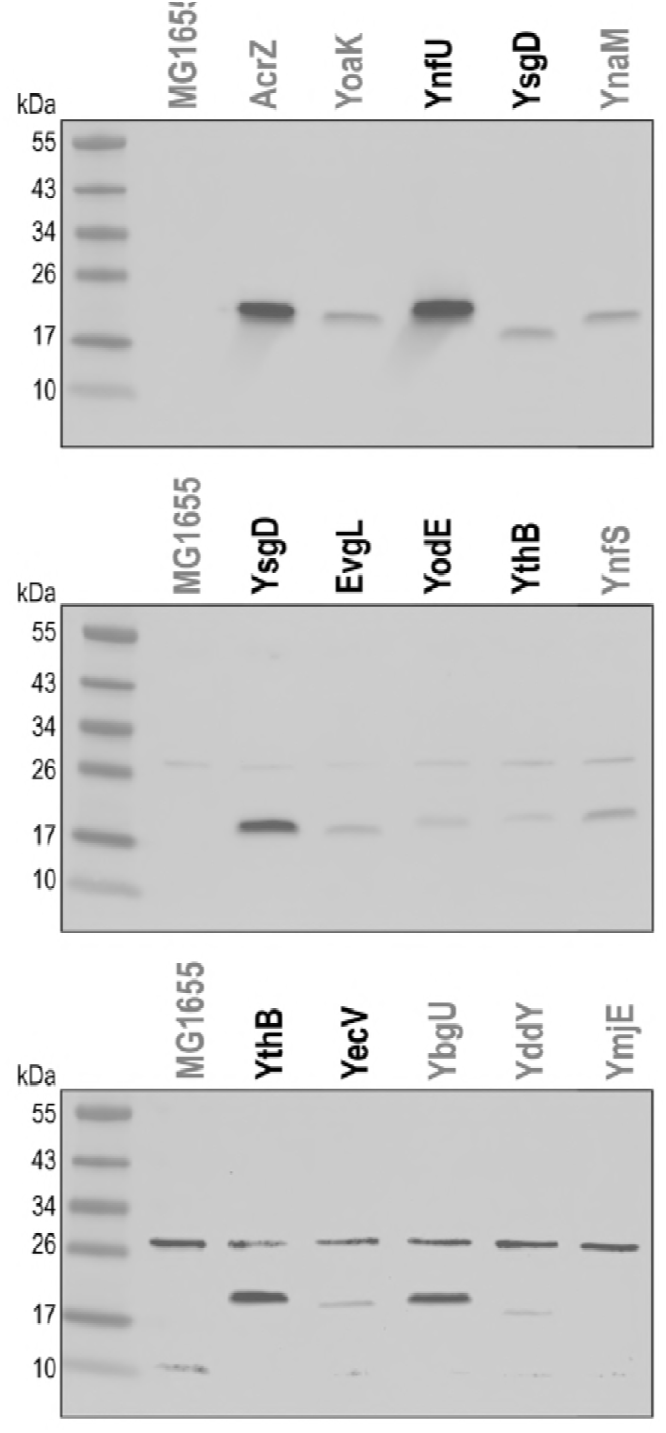
Observed small protein levels span several orders of magnitude. Stationary phase samples grown in LB from (Fig. 3) were compared to each other and to similarly-prepared samples of previously-detected small proteins (gray) with the same chromosomal tag (17, 37). Immunoblot analysis was carried out as for Fig. 3 with MG1655 as a negative control. All samples are in the MG1655 background and equally loaded, except for AcrZ, where the sample was diluted 1:5.

We also compared the levels of the new small proteins to five (YnaM, YnfS, YgbU, YddY and YmjE) of the 36 small proteins identified more recently (17). Three of the tested proteins (YnaM, YnfS and YbgU) are observed at levels comparable to most of the newly-identified small proteins, while two (YddY and YmjE) are more comparable to the least-abundant small proteins identified in this study (Fig. 4). It is interesting to note that YnaM, which had no ribosome density at start codons in the presence of retapamulin or Onc112, was detected at higher levels than most of the newly-detected small proteins, while YnfS, which has strong start peaks in both antibiotic-treated samples, was detected at lower levels.

### Some small proteins are encoded antisense to genes encoding expressed proteins

Given that antisense transcription in bacteria frequently is a means of gene silencing (reviewed in (38, 39)), we were surprised to note that eight of the newly detected proteins are encoded antisense to annotated protein-coding genes (Table 1). Additionally, one predicted smORF, *yoaM*, could not be tagged as it is found antisense to the operon of the essential *nrdA* and *nrdB* genes (encoding ribonucleoside-diphosphate reductase 1) (Fig. 5a). To test for expression of YoaM, we generated a translational fusion at the *lacZ* locus. Consistent with translation of this antisense-encoded small protein, we detect higher β-galactosidase expression for the *yoaM-lacZ* fusion than for an out-of-frame control fusion (Fig. 5e). Given that a clear transcriptional start was noted 174 nucleotides upstream of the YoaM start codon (40), it is possible that the synthesis of this protein is under post-transcriptional regulation.

**TABLE 1.**
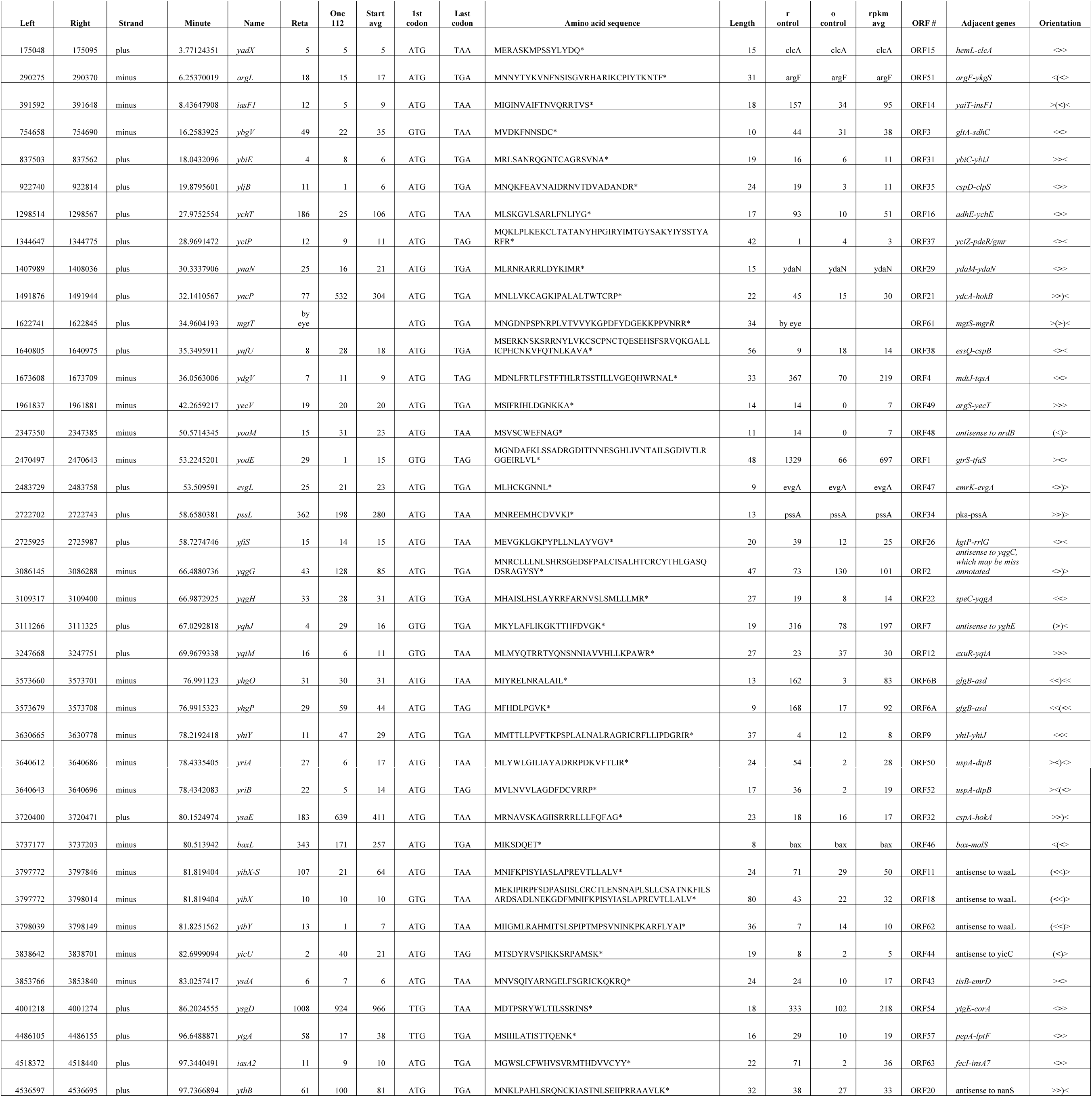
New small proteins detected. For each gene, we list the genomic co-ordinates (left and right), sense strand, position in the chromosome (in minutes, the map coordinates are based on the latest annotated version of the MG1655 genome in GenBank dated 24-SEP-2018, with a genome size of 4641652 bp), systematic gene name, start peak intensity in retapamulin- or Onc112-treated samples (in rpm), start and stop codons, peptide sequence and length, level of normal ribosome profiling density in untreated controls (in rpkm), candidate ORF number, and the orientation and names of neighboring genes.

**FIG 5.**
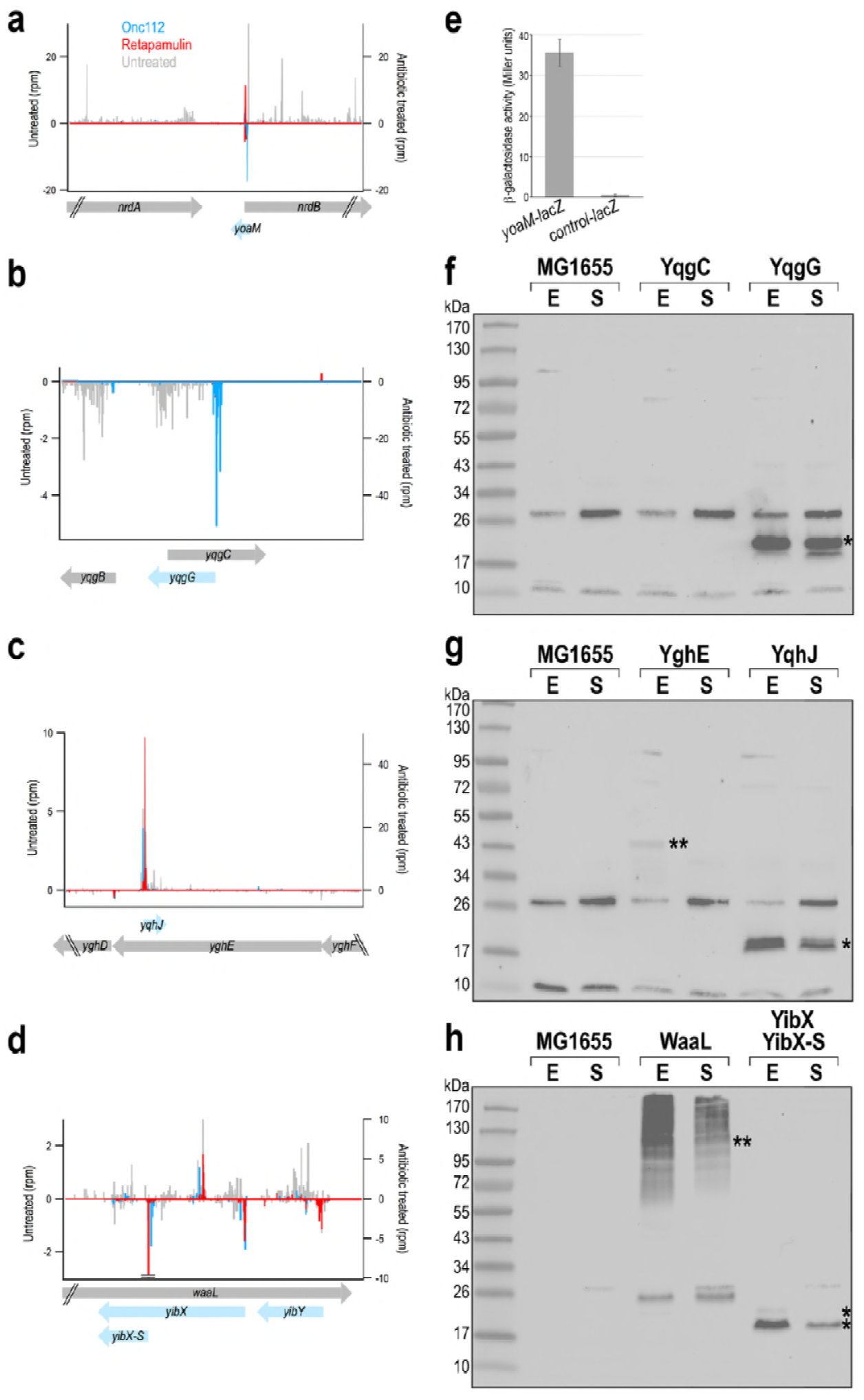
Novel smORFs (blue) are transcribed antisense to known genes (grey). The gene organization for the *nrdB-yoaM* (a), *yqgC-yqgG* (b), *yghE-yqhJ* (c) and *waaL-yibX-yibY* (d) loci. β-galactosidase activity was assayed for cells carrying chromosomal fusions of the 5’-UTR and initial codons of *yoaM* fused to *lacZ* as well as out-of-frame control fusion (e), which were grown in rich media (LB) with 0.2% arabinose. Protein levels for chromosomally SPA-tagged *yqgC* and *yqgG* (f), *yghE* and *yqhJ* (g) and *waaL* and *yibX* (h) genes. Gel samples were prepared from MG1655 strains grown to exponential (E) and stationary (S) phase in LB. Immunoblot analysis was carried out as for Fig. 3 with MG1655 as a negative control. Bands corresponding to smORFs are marked with *, and bands corresponding to antisense-encoded larger proteins are marked with **.

We wanted to determine whether annotated proteins and the newly-identified small proteins encoded by transcripts on opposing strands are both synthesized. We therefore introduced chromosomal tags upstream of the stop codons of the previously-annotated genes *yqgC* (antisense to *yqgG*) (Fig. 5b), *yghE* (antisense to *yqhJ*) (Fig. 5c) and *waaL* (antisense to *yibX* and *yibY*) (Fig. 5d). The *yqgC* gene (a protein of unknown function) does not have any associated ribosome density in either treated or untreated cells and the corresponding tagged protein is not observed under these conditions (Fig. 5f). YghE (another protein of unknown function), while detected, appears to be present at lower levels than YqhJ (Fig. 5g), consistent with its low levels of normal ribosome density (not visible at the scale used in Fig. 5c). WaaL (an O-antigen ligase) was clearly detected under the same growth conditions as YibX and YibX-S (Fig. 5h). We suggest the appearance of a smear for WaaL may be due to bound oligosaccharide substrates. In general, our results confirm that proteins can be encoded by both strands of the same region of DNA and expressed under the same growth conditions.

### YibX is translated as two isoforms

The *yibX* gene was also interesting as the profiling data suggested translation could initiate from two different start codons. While most bacterial ORFs encode a single protein, there are some examples where different isoforms of the same protein are generated by different translation starts in the same frame, as has been found for the *E. coli* proteins ClpB, IF-2, and MrcB (41-43). Frequently, the longer polypeptide is expressed at higher levels than the shorter isoform. A broad peak near the start codon for the ribosome profiling data suggests that several small proteins are potentially translated as different isoforms. Although most of the potential isoforms vary by only a few codons and would be indistinguishable on immunoblots, the YibX alternative start sites lead to proteins of substantially different sizes. The stronger signal corresponds to the 24-aa YibX-S protein, while a second signal at a GTG codon upstream and in frame with YibX-S yields an 80-aa protein adding ∼6.1 kDa (Fig. 5h). Both bands are detected in Fig. 3 and Fig. 5, but contrary to other known primary isoforms, the 80-amino acid protein is detected at lower levels than the shorter isoform. A second example of possible isoforms is YqhJ, which shows two bands in Fig 5g. YqhJ initiates at a GTG codon and is 19 residues long; initiation at a downstream TTG codon would yield a 13-residue protein (Table S3). Ribosome density in retapamulin and Onc112 treated samples is consistent with both of these initiation sites being used (data not shown).

### Multiple smORFs are encoded by different, overlapping frames

There are a growing number of bacterial examples where more than one protein is encoded in the same region in different frames, as has been found for *rzoD* encoded within *rzpD*, which are homologous to the rz/rz1 lysis cassette of bacteriophage λ (44, 45). A similar gene arrangement of nested start codons and substantial overlap is also found for two sets of newly-identified small proteins: YhgO/YhgP (Fig. 6a) and YriA/YriB (Fig. 6b). Additionally, the smORFs encoding two other new proteins, YbgV and MgtT, overlap the 3’-ends of the previously identified smORFs *ybgU* (Fig. 6c) and *mgtS* (Fig. 6d), respectively. We sought to compare the levels of the paired small proteins under the same conditions by assaying cells with one or the other smORF tagged (Fig. 6e-6h). Although there generally appears to be limited correlation between ribosome density and observed protein levels, for each of these pairs, the small protein corresponding to the smORF with the higher ribosome density with either retapamulin or Onc112 treatment (YhgP, YriB, YbgV and MgtS) was present at higher levels. Perhaps there is a better correlation between ribosome density and observed protein levels for co-transcribed genes.

**FIG 6.**
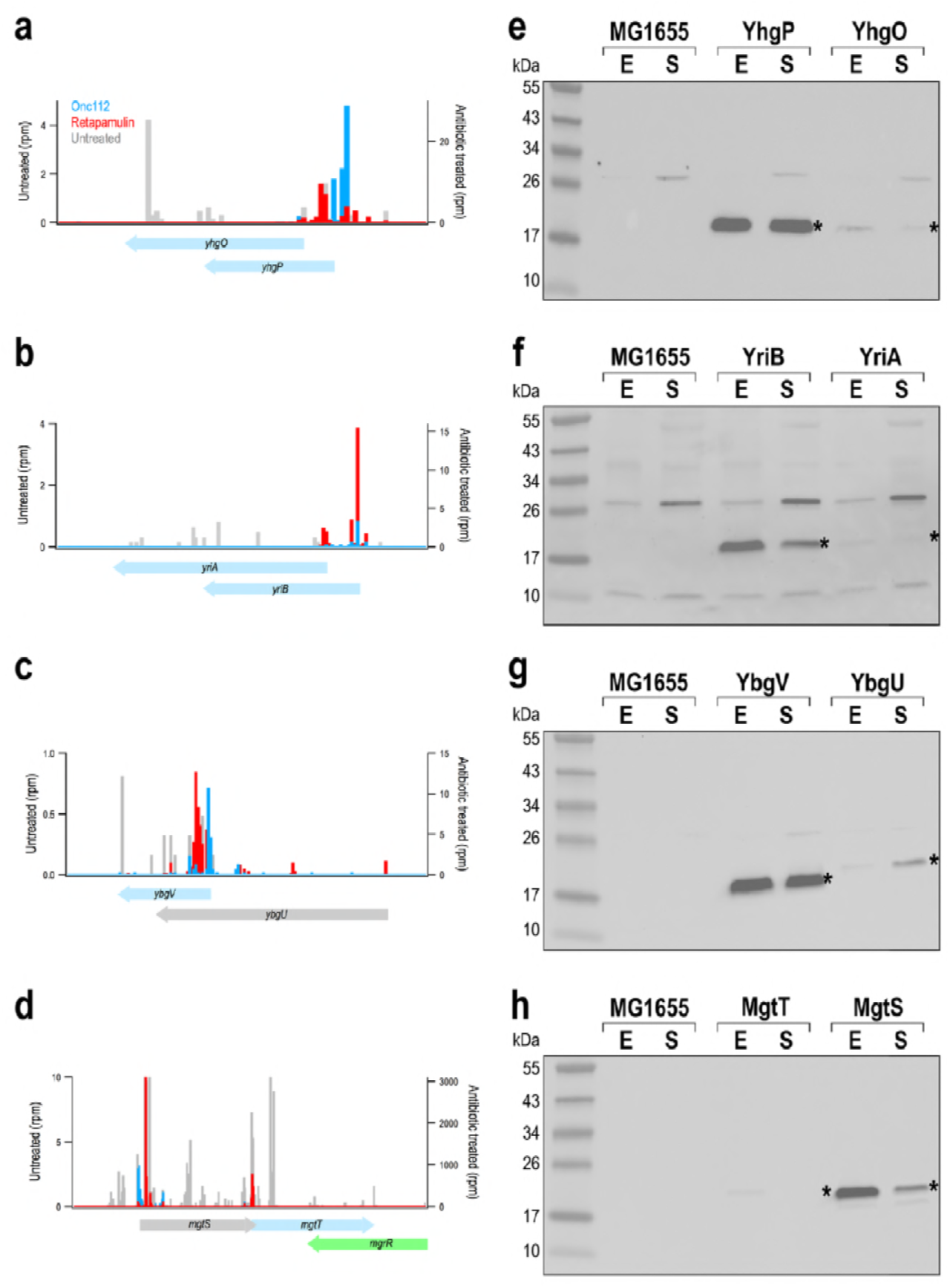
smORFs are organized in complex gene arrangements. Gene organization for *yhgO/yhgP* (a), *yriA/yriB* (b), *ybgU/ybgV* (c) and *mgtS/mgtT* (d), with previously-identified small protein genes in gray, newly-identified small protein genes in blue and small RNA gene *mgrR* in green. (b) Levels of corresponding proteins (e-h). Gel samples were prepared from MG1655 strains grown to exponential (E) and stationary (S) phase in LB. Immunoblot analysis was carried out as for Fig. 3 with MG1655 as a negative control. Bands corresponding to smORFs are marked with *.

### smORFs overlap the 5′ ends of larger protein coding genes

The genes of three new small proteins detected by immunoblot analysis (Fig. 3) were found to overlap the 5′end of annotated larger genes in a different frame: *baxL*-*baxA, evgL*-*evgA*, and *argL*-*argF*. Two additional smORFs predicted by ribosome profiling, *ORF33* and *pssL*, also overlap the 5′ end of the neighboring gene in a different frame, but we were unable to SPA-tag these predicted proteins given the downstream genes, *accD* (acetyl-CoA carboxyltransferase subunit β) (Fig. 7a) and *pssA* (phosphatidylserine synthase) (Fig. 7b), are essential. To investigate the expression of ORF33 and PssL, the 5’-UTR and the first few codons of the smORFs were translationally fused to *lacZ* on the chromosome (46). While there was no measurable β-galactosidase activity for the *ORF33-lacZ* fusion (Fig. 7f), there was clear expression of the *pssL-lacZ* fusion, which was diminished by the introduction of a stop codon at the start codon position (Fig. 7g). These results indicate that although we could not construct a *pssL-SPA* fusion at the endogenous location of the genome, the protein is translated.

**FIG 7.**
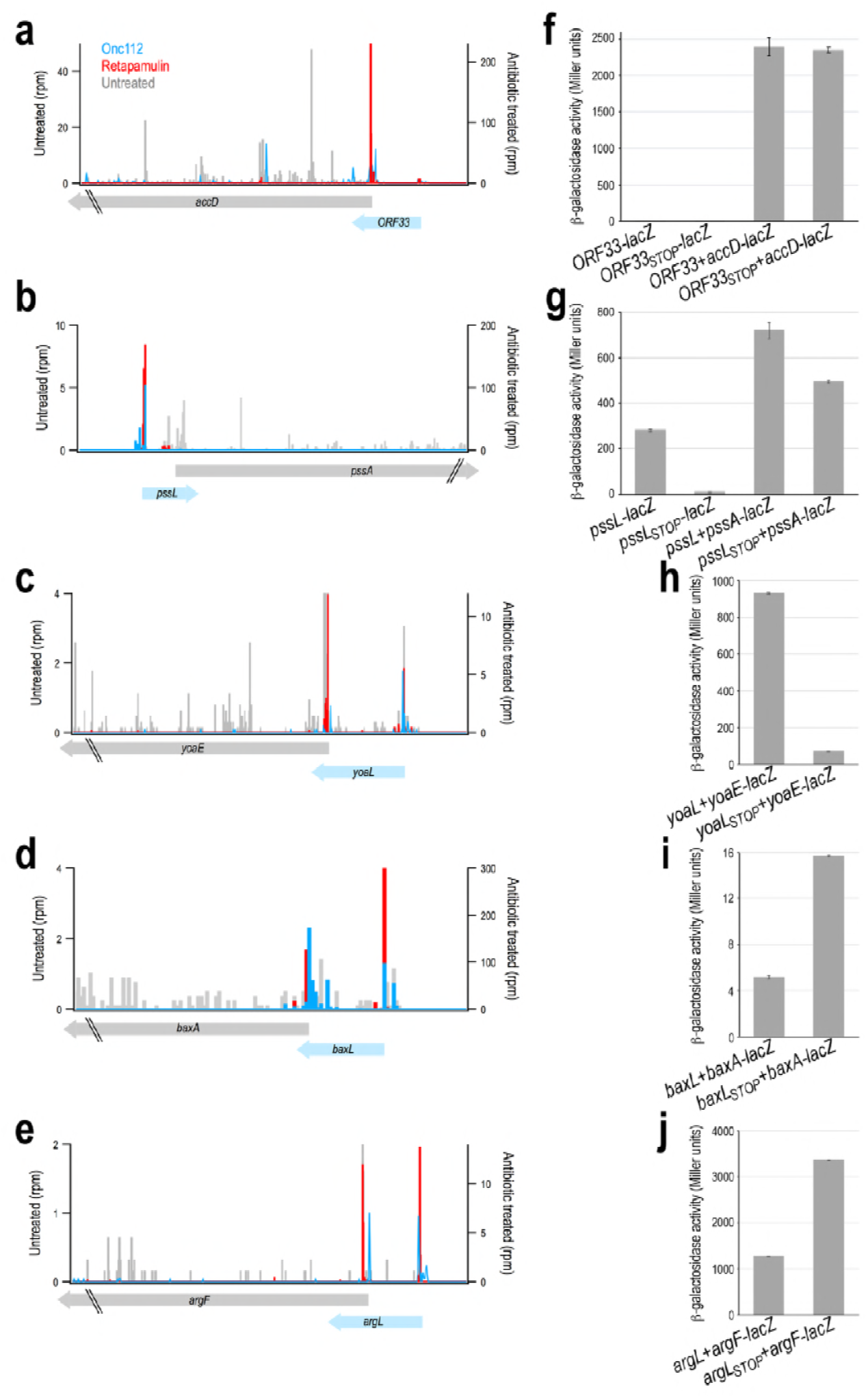
smORFs function as leader peptides to regulate translation of downstream genes. (a-e) Organization of smORFs (blue) in 5’-UTRs of known genes (gray). β-galactosidase activity was assayed for cells carrying chromosomal fusions of the 5’-UTR and initial codons of *ORF33* (f) and *pssL* (g) fused to *lacZ*. β-galactosidase activity was assayed for cells carrying *lacZ* chromosomal fusions to the 5’-UTR and initial codons of the downstream gene with a wild type start codon for the upstream smORF or with a stop codon replacing the start (f-j). For all β-galactosidase assays, cells were grown in rich media (LB) with 0.2% arabinose before enzyme activity was measured.

### Role of smORFs regulating expression of larger protein encoded downstream

Given other examples where smORFs overlapping downstream genes serve as leader peptides involved in modulating the translation of the larger gene ((47), reviewed in (48)), we next sought to investigate whether translation of the smORFs overlapping larger ORFs described above affects translation of the downstream ORF. To test this, the entire 5′ UTR including the smORF together with the first codons of the downstream gene was fused to *lacZ* at the endogenous *lacZ* locus. We also generated a second version of these constructs by introducing amber or ochre stop codons into the smORF as a replacement for the start codon. If translation of the two ORFs is coupled, the stop codon, which blocks the expression of the upstream smORF, should impact translation of the downstream gene. In the case of the *ORF33-accD* pair, for which we did not see any expression of ORF33, the stop codon had no impact on *accD-lacZ* expression (Fig. 7f). In contrast, introduction of a stop codon into *pssL* led to a 30% decrease in the expression of the *pssA-lacZ* fusion (Fig. 7g), while introduction of a stop codon into *yoaL*, a recently identified smORF (17), led to strongly decreased expression of *yoaE-lacZ* (Fig. 7h). An increase in the expression of the downstream gene is observed when stop codons are introduced into *baxL* (Fig. 7i) and *argL* (Fig. 7j). Together these results indicate that translation of these upstream smORFs may be playing a regulatory role.

## DISCUSSION

Fundamentally, the challenge of identifying expressed small proteins stems from the great number of putative smORFs, with ∼161,000 possible smORFs in intergenic regions of *E. coli* alone. The key question is how to best identify and validate candidate smORFs in a manner that prevents the annotation of uncorroborated genes. Rather than relying solely on bioinformatics approaches, as has been done previously, we demonstrated that an approach that utilizes multiple ribosome profiling datasets can identify translated smORFs with a high degree of accuracy. The expression of 36 of these smORFs was verified by immunoblot analysis of the chromosomally-tagged genes, and the expression of two other genes that could not be tagged at the endogenous loci was observed as chromosomal *lacZ* fusions. We noted a number of interesting gene arrangements including small proteins encoded on the strand opposite larger, annotated proteins, as well as smORFs in the 5′-UTR of known genes.

### Limitations of approach

While we were able to identify many new small proteins, we are cognizant of some limitations. One important caveat is that start codon peak intensity in profiling experiments is not a truly quantitative measure of initiation rates, given that reads at a single site are prone to sequence-specific artifacts (49). Examination of the profiling data of previously-identified smORFs illustrates this limitation, as there is not a strong correlation between ribosome density in the presence of the initiation complex inhibitors and the band intensity observed by immunoblot analysis. While the degradation of some tagged small proteins may explain ribosome density without corresponding protein bands, other smORFs yield strong bands without any sequencing reads in the profiling experiments. Determining the factors that contribute to the perceived mismatch between ribosome density and observed protein levels would allow for a more accurate prediction of expression. It also must be considered that, although only occurring for a short duration, treatment with retapamulin or Onc112 represents a stress upon the bacteria that can cause changes in the expression profile.

One other major limitation regarding the general application of this approach is that the microbes must be susceptible to these initiation complex inhibitors. Retapamulin is a member of the pleuromutilin class of antibiotics that show activity against a broad spectrum of gram-positive bacteria, though some derivatives show activity against gram-negative bacteria as well (50, 51). To increase susceptibility to retapamulin, the group of Vázquez-Laslop and Mankin (35) used a *tolC* mutant strain of *E. coli*, an approach that may need to be employed in other bacteria. Onc112, a member of the PrAMP family of peptide antibiotics, is actively transported into gram-negative bacteria by proteins such as the SbmA transporter (52). It may be possible to extend the range of compounds like Onc112 by exogenous expression of transporters such as SbmA in bacteria that otherwise lack them.

### Advantages of approach

Despite the possible limitations, the ability to identify start codons through ribosome profiling with inhibitors is a powerful approach with broad applications. As shown here, translated smORFs are more prevalent than previously believed and are found in contexts that would be difficult to distinguish by other methods, including bioinformatics approaches that have been successfully employed previously (16, 17). While traditional ribosome profiling can guide the prediction (53, 54) or, in conjunction with experiments to verify protein synthesis, even support the annotation of intergenic smORFs in bacteria (27), ribosome profiling with stalled initiation complexes allows for new ORFs to be located in contexts that are generally-ignored, including within or overlapping other genes as shown here and by the group of Vázquez-Laslop and Mankin (35). These new, internal altORFs may represent a new class of functional and regulatory proteins that comprise an ever-expanding proteome.

Interestingly, we noted relatively poor overlap between our predicted smORFs and those reported in the other ribosome profiling studies (27, 31) suggesting that many small proteins remain to be discovered. Of the 328 smORFs predicted by Mori and co-workers in intergenic regions in *E. coli* based on ribosome enrichment at start codons after treatment with tetracycline, only 20 overlap with our list of 68 likely candidates (Table S3). The fact that retapamulin and Onc112 are more specific than tetracycline for newly initiated ribosomes, providing higher resolution for start codon identification, may partially explain the limited overlap with our predicted smORFs. We also looked for overlap between our 68 likely candidates and the 130 smORFs predicted in *Salmonella enterica* in a recent study using traditional ribosome profiling (27). Only one exact match and three close matches were found between these closely related species.

In addition to facilitating the identification of new smORFs, the profiling data with inhibitors provide valuable information about known ORFs and suggest the need to reannotate some genes (Table S1). One example is the smORF *ymiA*, which is annotated both as beginning with MLISDGDYMRLAMPSGNQEP (55) and as beginning with the third methionine at MPSGNQEP (16), but likely initiates with MRLAMP (Fig. S3). Another example is the *ymdG* protein. Although it is annotated as 40 residues, our data show that a later start codon is used and that the smORF is only eight codons long (Fig. S3). Finally, *yoaL*, which was herein examined for function as a leader peptide (Fig. 7), was originally annotated as initiating on a methionine 13 codons upstream of its likely start site (17) (Fig. S3).

### Small protein function

Many of the small proteins are expected to have functions that involve the binding to other, larger proteins. However, the primary structures of the small proteins are often too short for bioinformatics tools to identify motifs or domains that may offer insights into their functions in the cell. Of the newly-identified small proteins, only YnfU, which is encoded within the Qin prophage region of the *E. coli* genome, had an identifiable motif. The protein contains a pair of zinc knuckles, a motif with two copies of the CPXC sequence that together chelate a zinc ion (reviewed in (56)). Homology modelling of YnfU using PSIPRED (57) also revealed a moderate match to the zinc-binding domain of PA0128, a protein of unknown function from *Pseudomonas aeruginosa*.

Although motif identification is often not available for smORFs, multiple previously-identified proteins were predicted to contain transmembrane helices and were later experimentally verified to localize to the cellular membrane (16). When we examined the sequences of the new smORFs using the Phobius or ExPASy TMpred algorithms (58, 59), none of the newly-identified proteins were predicted to contain transmembrane helices. This analysis shows that the skew toward hydrophobic α-helices overall is not as strong as observed for the first small *E. coli* proteins identified (16). In general, the next challenge will be to determine functions for the large numbers of newly-identified proteins.

Four new smORFs and one previously annotated smORF were examined for possible roles as leader peptides, as these small protein genes overlap the downstream coding sequences of larger proteins in alternate frames (Fig. 7). For each of the expressed genes, either an increase or decrease in the translation of the downstream gene was observed when the upstream smORF was not translated. For genes where expression decreases, this drop may stem from a loss of translational coupling from the upstream gene, while for genes with improved expression, translation of the smORF may impede translation of the downstream gene. It is interesting to note that a mutation (*pssR1*) that leads to increased expression of *pssA* mapped to the anti-Shine-Dalgarno sequence of the 16S rRNA encoded by *rrnC* (60). Further characterization will be required to distinguish smORFs that are simply translated in operons versus those that specifically serve to control the translation of downstream genes as well as to elucidate the regulatory mechanisms.

### Complex gene organization

Beyond the expanded presence of smORFs as possible upstream leaders of other genes, our analysis also pointed to other forms of complex gene organization. We found several smORFs that overlap other new or known smORFs. We also discovered small proteins encoded antisense to larger proteins, as well as at least one small protein that is translated as two isoforms. We hypothesize the pairs of bacterial genes encoded in overlapping regions have related functions.

Since we think we have not yet identified the complete set of small protein genes, we suggest that antisense genes and translational regulation by upstream smORFs may be far more prevalent than currently thought. Full annotation of translated regions of the chromosome will be required to obtain a more comprehensive picture of cellular regulation. Additionally, more complete annotation will provide a better understanding of the roles of the many seemingly orphan transcription start sites observed in transcriptome data (40). The use of ribosome profiling with initiation complex inhibitors revealed 38 new protein-coding genes in *E. coli*, an organism already known to express nearly 100 small proteins. For less well characterized bacteria, the ability to define the small proteome accurately and in an unbiased manner opens new doors to uncovering the regulation that allows the growth and survival of these organisms.

## MATERIALS AND METHODS

### Onc112 ribosome profiling

A culture of *E. coli* MG1655 was grown overnight at 37°C in MOPS EZ Rich Defined media (Teknova) with 0.2% glucose, diluted 1:100 into 150 mL of fresh media, and grown to OD_600_ = 0.3. The culture was treated with 50 µM Onc112 for 10 min and harvested by rapid filtration and freezing in liquid nitrogen. Ribosome profiling libraries were prepared and sequenced as described (61) except that the lysis buffer contained 1 M NaCl to arrest translation instead of chloramphenicol. Following lysis and clarification, 25 AU of RNA in the lysate was pelleted over a 1 mL sucrose cushion (20 mM Tris pH 7.5, 500 mM NH_4_Cl, 0.5 mM EDTA, 1.1 M sucrose) using a TLA 100.3 rotor at 65,000 rpm for 2 h. Pellets were resuspended in 200 µL of the standard lysis buffer and the RNA was digested with MNase following the standard protocol.

### Analysis of ribosome profiling data

Raw reads were filtered and trimmed using Skewer v0.2.2. Reads were mapped uniquely to the MG1655 genome NC_000913.3 (allowing two mismatches) using Bowtie v 0.12.7 after reads mapping to tRNA and rRNA were discarded. Ribosome density was assigned to the 3’-end of reads. We identified novel open reading frames 8 sense codons or longer starting with ATG, GTG, or TTG codons at least 18 nt away (on either side) from any annotated genes. For each candidate, the ribosome density in retapamulin or Onc112 treated samples was summed 0-18 nt downstream of the first nt in the start codon to calculate the initiation peak intensity. We also calculated rpkm values for normal ribosome profiling data for each candidate smORF unless any part of it comes within 15 nt of an annotated gene. Each of these candidates and their scores are reported in Table S2. The retapamulin data can be found at GSE122129 and the Onc112 data can be found at GSE123675.

### Strain construction

All strains generated for this study are listed in Table S4 together with the sequences of the oligonucleotides used to construct the strains. smORFs were tagged on the chromosome following published procedures (62). In short, an SPA-kan cassette was inserted at the C-terminal end of each ORF using the λ Red recombination system in *E. coli* NM400 and moved into *E. coli* MG1655 by P1 transduction. All insertions were verified by sequencing.

Construction of the *lacZ* reporter strains followed published procedure (46). In brief, DNA including the 5’-UTR and several codons of each ORF, along with flanking homology regions, were transformed into *E. coli* PM1205, which utilizes the λ Red-mediated recombination system, and selected for sucrose resistance. All insertions were verified by sequencing.

### Immunoblot analysis

For all expression experiments, Luria broth (LB) was inoculated 1:200 with overnight culture of various strains and grown at 37°C. One mL of culture was taken during exponential growth (two h post inoculation, OD_600_ = 0.5-0.7) and during stationary phase (3.5 h post inoculation, OD_600_ = 2.5-3). To normalize for total cell (number/density/count), the cell pellet collected for each sample was resuspended according to the OD_600_. Samples were analyzed on SDS-PAGE, transferred to nitrocellulose membranes, and blotted using anti-FLAG(M2)-HRP (Sigma).

### Assays of β-galactosidase activity

For all experiments, LB with 0.2% arabinose was inoculated 1:200 with overnight culture of PM1205 strains carrying various *lacZ-*fusions. These cultures were grown at 37°C for 2.25 h (OD600 = 0.75-1.0). Culture was added directly to Z-buffer in 1.5 ml microcentrifuge tubes. SDS (0.00184%) and chloroform (3.5% v/v) were added, and samples were vortexed for 30 seconds. The samples were incubated at 28°C for 15 min before the addition of ortho-nitrophenyl-β-galactoside (ONPG, 0.875 mg/ml). Incubation at 28°C continued until a visible color change occurred, at which time sodium carbonate (353 mM) was added to quench the reaction. All reactions were quenched by 75 min, even if no color change was observed. Samples were centrifuged at maximum speed in a table-top microcentrifuge (∼21000 x g) for 2 min. 1 ml of supernatant was used to measure the absorbance at 550 nm and 420 nm. Miller Units were calculated using the established formula (63).

## ACKNOWLEDGMENTS

The authors thank Alexander Mankin and Nora Vázquez-Laslop for sharing the retapamulin-treated ribosome profiling data prior to publication and Matthew Hemm for sharing strains prior to publication. We also thank A. Mankin, N. Vázquez-Laslop, P. Adams and M. Wu Orr for comments on the manuscript. Work in the G.S. laboratory was supported by the Intramural Research Program of the *Eunice Kennedy Shriver* National Institute of Child Health and Human Development. Work in the A.B. laboratory was supported by grants from the National Institute of General Medical Sciences (GM110113 and GM105816).

## REFERENCES

1. Storz G, Wolf YI, Ramamurthi KS. 2014. Small proteins can no longer be ignored. Annu Rev Biochem 83:753–777.

2. Andrews SJ, Rothnagel JA. 2014. Emerging evidence for functional peptides encoded by short open reading frames. Nat Rev Genet 15:193–204.

3. Saghatelian A, Couso JP. 2015. Discovery and characterization of smORF-encoded bioactive polypeptides. Nat Chem Biol 11:909–916.

4. Hobbs EC, Yin X, Paul BJ, Astarita JL, Storz G. 2012. Conserved small protein associates with the multidrug efflux pump AcrB and differentially affects antibiotic resistance. Proc Natl Acad Sci USA 109:16696–16701.

5. Wang H, Yin X, Wu Orr M, Dambach M, Curtis R, Storz G. 2017. Increasing intracellular magnesium levels with the 31-amino acid MgtS protein. Proc Natl Acad Sci USA 114:5689–5694.

6. Yin X, Wu Orr M, Wang H, Hobbs EC, Shabalina SA, Storz G. 2018. The small protein MgtS and small RNA MgrR modulate the PitA phosphate symporter to boost intracellular magnesium levels. Mol Microbiol In press.

7. Bal NC, Maurya SK, Sopariwala DH, Sahoo SK, Gupta SC, Shaikh SA, Pant M, Rowland LA, Bombardier E, Goonasekera SA, Tupling AR, Molkentin JD, Periasamy M. 2012. Sarcolipin is a newly identified regulator of muscle-based thermogenesis in mammals. Nat Med 18:1575–1579.

8. Anderson DM, Anderson KM, Chang CL, Makarewich CA, Nelson BR, McAnally JR, Kasaragod P, Shelton JM, Liou J, Bassel-Duby R, Olson EN. 2015. A micropeptide encoded by a putative long noncoding RNA regulates muscle performance. Cell 160:595–606.

9. Goffeau A, Barrell BG, Bussey H, Davis RW, Dujon B, Feldmann H, Galibert F, Hoheisel JD, Jacq C, Johnston M, Louis EJ, Mewes HW, Murakami Y, Philippsen P, Tettelin H, Oliver SG. 1996. Life with 6000 genes. Science 274:546–567.

10. Burge CB, Karlin S. 1998. Finding the genes in genomic DNA. Curr Opin Struct Biol 8:346–354.

11. Congdon RW, Muth GW, Splittgerber AG. 1993. The binding interaction of Coomassie blue with proteins. Anal Biochem 213:407–413.

12. Scopes RK. 1994. Protein Purification Principles and Practice, 3 ed. Springer-Verlag, New York, NY.

13. Burgess R, Deutscher M (ed). 2009. Guide to Protein Purification. Academic Press, San Diego, CA.

14. Slavoff SA, Mitchell AJ, Schwaid AG, Cabili MN, Ma J, Levin JZ, Karger AD, Budnik BA, Rinn JL, Saghatelian A. 2013. Peptidomic discovery of short open reading frame-encoded peptides in human cells. Nat Chem Biol 9:59–64.

15. Pueyo JI, Magny EG, Couso JP. 2016. New peptides under the s(ORF)ace of the genome. Trends Biochem Sci 41:665–678.

16. Hemm MR, Paul BJ, Schneider TD, Storz G, Rudd KE. 2008. Small membrane proteins found by comparative genomics and ribosome binding site models. Mol Microbiol 70:1487–501.

17. VanOrsdel CE, Kelly JP, Burke BN, Lein CD, Oufiero CE, Sanchez JF, Wimmers LE, Hearn DJ, Abuikhdair FJ, Barnhart KR, Duley ML, Ernst SEG, Kenerson BA, Serafin AJ, Hemm MR. 2018. Identifying new small proteins in *Escherichia coli*. Proteomics 18:e1700064.

18. Ladoukakis E, Pereira V, Magny EG, Eyre-Walker A, Couso JP. 2011. Hundreds of putatively functional small open reading frames in *Drosophila*. Genome Biol 12:R118.

19. Kessler MM, Zeng Q, Hogan S, Cook R, Morales AJ, Cottarel G. 2003. Systematic discovery of new genes in the *Saccharomyces cerevisiae* genome. Genome Res 13:264–271.

20. Landry CR, Zhong X, Nielly-Thibault L, Roucou X. 2015. Found in translation: functions and evolution of a recently discovered alternative proteome. Curr Opin Microbiol 32:74–80.

21. Mouilleron H, Delcourt V, Roucou X. 2016. Death of a dogma: eukaryotic mRNAs can code for more than one protein. Nucleic Acids Res 44:14–23.

22. Vanderperre B, Lucier JF, Bissonnette C, Motard J, Tremblay G, Vanderperre S, Wisztorski M, Salzet M, Boisvert FM, Roucou X. 2013. Direct detection of alternative open reading frames translation products in human significantly expands the proteome. PLoS ONE 8:e70698.

23. Hellens RP, Brown CM, Chisnall MA, Waterhouse PM, Macknight RC. 2016. The emerging world of small ORFs. Trends Plant Sci 21:317–328.

24. Neuhaus K, Landstorfer R, Simon S, Schober S, Wright PR, Smith C, Backofen R, Wecko R, Keim DA, Scherer S. 2017. Differentiation of ncRNAs from small mRNAs in *Escherichia coli* O157:H7 EDL933 (EHEC) by combined RNAseq and RIBOseq -*ryhB* encodes the regulatory RNA RyhB and a peptide, RyhP. BMC Genomics 18:216.

25. Bazzini AA, Johnstone TG, Christiano R, Mackowiak SD, Obermayer B, Fleming ES, Vejnar CE, Lee MT, Rajewsky N, Walther TC, Giraldez AJ. 2014. Identification of small ORFs in vertebrates using ribosome footprinting and evolutionary conservation. EMBO J 33:981–993.

26. Aspden JL, Eyre-Walker YC, Phillips RJ, Amin U, Mumtaz MA, Brocard M, Couso JP. 2014. Extensive translation of small Open Reading Frames revealed by Poly-Ribo-Seq. Elife 3:e03528.

27. Baek J, Lee J, Yoon K, Lee H. 2017. Identification of unannotated small genes in *Salmonella*. G3 7:983–989.

28. Guttman M, Russell P, Ingolia NT, Weissman JS, Lander ES. 2013. Ribosome profiling provides evidence that large noncoding RNAs do not encode proteins. Cell 154:240–251.

29. Ingolia NT, Lareau LF, Weissman JS. 2011. Ribosome profiling of mouse embryonic stem cells reveals the complexity and dynamics of mammalian proteomes. Cell 47:789–802.

30. Lee S, Liu B, Lee S, Huang SX, Shen B, Qian SB. 2012. Global mapping of translation initiation sites in mammalian cells at single-nucleotide resolution. Proc Natl Acad Sci USA 109:E2424–E2432.

31. Nakahigashi K, Takai Y, Kimura M, Abe N, Nakayashiki T, Shiwa Y, Yoshikawa H, Wanner BL, Ishihama Y, Mori H. 2016. Comprehensive identification of translation start sites by tetracycline-inhibited ribosome profiling. DNA Res 23:193–201.

32. Seefeldt AC, Nguyen F, Antunes S, Pérébaskine N, Graf M, Arenz S, Inampudi KK, Douat C, Guichard G, Wilson DN, Innis CA. 2015. The proline-rich antimicrobial peptide Onc112 inhibits translation by blocking and destabilizing the initiation complex. Nat Struct Mol Biol 22:470–475.

33. Gagnon MG, Roy RN, Lomakin IB, Florin T, Mankin AS, Steitz TA. 2016. Structures of proline-rich peptides bound to the ribosome reveal a common mechanism of protein synthesis inhibition. Nucleic Acids Res 44:2439–2450.

34. Dornhelm P, Högenauer G. 1978. The effects of tiamulin, a semisynthetic pleuromutilin derivative, on bacterial polypeptide chain initiation. Eur J Biochem 91:465–473.

35. Meydan S, Marks J, Klepacki D, Sharma V, Baranov P, Firth A, Margus T, Kefi A, Vázquez-Laslop N, Mankin AS. 2018. Retapamulin-assisted Ribo-seq revels the alternative bacterial proteome. Mol Cell under review.

36. Roy RN, Lomakin IB, Gagnon MG, Steitz TA. 2015. The mechanism of inhibition of protein synthesis by the proline-rich peptide oncocin. Nat Struct Mol Biol 22:466–469.

37. Hemm MR, Paul BJ, Miranda-Rios J, Zhang A, Soltanzad N, Storz G. 2010. Small stress response proteins in *Escherichia coli*: proteins missed by classical proteomic studies. J Bacteriol 192:46–58.

38. Sesto N, Wurtzel O, Archambaud C, Sorek R, Cossart P. 2013. The excludon: a new concept in bacterial antisense RNA-mediated gene regulation. Nat Rev Microbiol 11:75–82.

39. Georg J, Hess WR. 2018. Widespread antisense transcription in prokaryotes. Microbiol Spectr 6:RWR–0029-2018.

40. Thomason MK, Bischler T, Eisenbart SK, Förstner KU, Zhang A, Herbig A, Nieselt K, Sharma CM, Storz G. 2015. Global transcriptional start site mapping using differential RNA sequencing reveals novel antisense RNAs in *Escherichia coli*. J Bacteriol 197:18–28.

41. Squires CL, Pedersen S, Ross BM, Squires C. 1991. ClpB is the *Escherichia coli* heat shock protein F84.1. J Bacteriol 173:4254–4262.

42. Sacerdot C, Dessen P, Hershey JW, Plumbridge JA, Grunberg-Manago M. 1984. Sequence of the initiation factor IF2 gene: unusual protein features and homologies with elongation factors. Proc Natl Acad Sci USA 81:7787–7791.

43. Ross TK, Achberger EC, Braymer HD. 1989. Nucleotide sequence of the McrB region of *Escherichia coli* K-12 and evidence for two independent translational initiation sites at the *mcrB* locus. J Bacteriol 171:1974–1981.

44. Zhang N, Young R. 1999. Complementation and characterization of the nested Rz and Rz1 reading frames in the genome of bacteriophage lambda. Mol Gen Genet 262:659–667.

45. Toba FA, Thompson MG, Campbell BR, Junker LM, Rueggeberg KG, Hay AG. 2011. Role of DLP12 lysis genes in *Escherichia coli* biofilm formation. Microbiology 157:1640–1650.

46. Mandin P, Gottesman S. 2009. A genetic approach for finding small RNAs regulators of genes of interest identifies RybC as regulating the DpiA/DpiB two-component system. Mol Microbiol 72:551–565.

47. Park H, McGibbon LC, Potts AH, Yakhnin H, Romeo T, Babitzke P. 2017. Translational repression of the RpoS antiadapter IraD by CsrA is mediated via translational coupling to a short upstream open reading frame. mBio 8:e01355–17.

48. Gollnick P, Babitzke P. 2002. Transcription attenuation. Biochim Biophys Acta 1577:240–250.

49. Gao X, Wan J, Liu B, Ma M, Shen B, Qian SB. 2015. Quantitative profiling of initiating ribosomes in vivo. Nat Methods 12:147–153.

50. Goethe O, Heuer A, Ma X, Wang Z, Herzon SB. 2018. Antibacterial properties and clinical potential of pleuromutilins. Nat Prod Rep In Press.

51. Novak R. 2011. Are pleuromutilin antibiotics finally fit for human use? Ann N Y Acad Sci 1241:71–81.

52. Mattiuzzo M, Bandiera A, Gennaro R, Benincasa M, Pacor S, Antcheva N, Scocchi M. 2007. Role of the *Escherichia coli* SbmA in the antimicrobial activity of proline-rich peptides. Mol Microbiol 66:151–163.

53. Hücker SM, Ardern Z, Goldberg T, Schafferhans A, Bernhofer M, Vestergaard G, Nelson CW, Schloter M, Rost B, Scherer S, Neuhaus K. 2017. Discovery of numerous novel small genes in the intergenic regions of the *Escherichia coli* O157:H7 Sakai genome. PLoS One 12:e0184119.

54. Friedman RC, Kalkhof S, Doppelt-Azeroual O, Mueller SA, Chovancová M, von Bergen M, Schwikowski B. 2017. Common and phylogenetically widespread coding for peptides by bacterial small RNAs. BMC Genomics 18:553.

55. Consortium. U. 2015. UniProt: a hub for protein information. Nucleic Acids Res 43:D204–212.

56. Krishna SS, Majumdar I, Grishin NV. 2003. Structural classification of zinc fingers: survey and summary. Nucleic Acids Res 31:532–550.

57. Lobley A, Sadowski MI, Jones DT. 2009. pGenTHREADER and pDomTHREADER: new methods for improved protein fold recognition and superfamily discrimination. Bioinformatics 25:1761–1767.

58. Ikeda M, Arai M, Okuno T, Shimizu T. 2003. TMPDB: a database of experimentally-characterized transmembrane topologies. Nucleic Acids Res 31:406–409.

59. Kall L, Krogh A, Sonnhammer EL. 2004. A combined transmembrane topology and signal peptide prediction method. J Mol Biol 338:1027–36.

60. Bartoli J, My L, Belmudes L, Couté Y, Viala JP, Bouveret E. 2017. The long hunt for *pssR*-looking for a phospholipid synthesis transcriptional regulator, finding the ribosome. J Bacteriol 199:e00202–17.

61. Woolstenhulme CJ, Guydosh NR, Green R, Buskirk AR. 2015. High-precision analysis of translational pausing by ribosome profiling in bacteria lacking EFP. Cell Rep 11:13–21.

62. Yu D, Ellis HM, Lee EC, Jenkins NA, Copeland NG, Court DL. 2000. An efficient recombination system for chromosome engineering in *Escherichia coli*. Proc Natl Acad Sci USA 97:5978–5983.

63. Miller JH. 1992. A Short Course in Bacterial Genetics: A Laboratory Manual and Handbook for Escherichia coli and Related Bacteria. Cold Spring Harbor Laboratory Press, Plainview, NY.

